# Single cell ATAC sequencing identifies sleepy macrophages during reciprocity of cytokines in *L.major* infection

**DOI:** 10.1101/2023.01.30.526191

**Authors:** Shweta Khandibharad, Komal Kharat, Shailza Singh

## Abstract

Hallmark for macrophages is their ability to possess plasticity which enables them to respond to changing microenvironment. *Leishmania* takes use of the phenotypic plasticity of macrophages to create favorable position for intracellular survival and persistent infection through regulatory cytokine like IL10. However, these effector cells can eliminate infection through modulation of critical cytokines such as IL12 and key players involved in its production. Using sophisticated tool of single cell ATAC sequencing we analyzed the regulatory axis of IL10 and IL12 in time dependent manner for *L.major* infection in macrophages. We recognized the cellular heterogeneity post infection with the regulators of IL10 and IL12 and observed a reciprocal relationship between them. Our prominent findings also exposed sleepy macrophages and its role in IL10 and IL12 reciprocity. To summarize, role of NFAT5 was vital in identification of sleepy macrophages which is a transient state where transcription factors controlled the epigenetic remodeling and expression of genes involved in pro-inflammatory cytokine production and recruitment of immune cells.

Graphical abstract summary of single cell ATAC sequencing for *L.major* infected RAW264.7 mouse macrophage cell line. (A) Reciprocal relationship identified in cell populations which is governed through chromatin remodeling. (B) Single cell ATAC pipeline for analysis of dataset (C) Identification of sleepy macrophages which may play potential role in reciprocity.

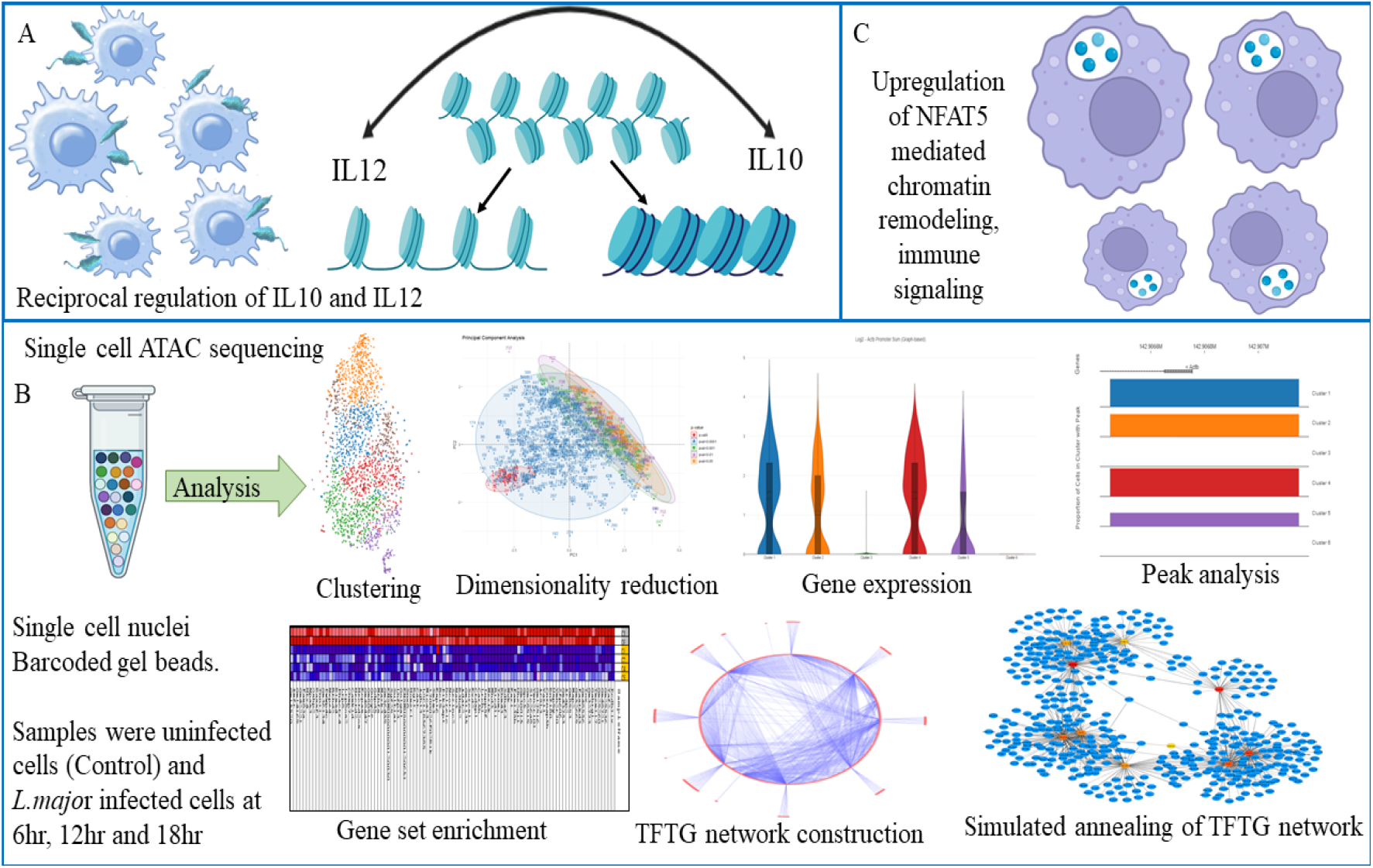

## Introduction

Effective immune functioning is influenced over the course of lifetime by niche of foreign entities through anthroponotic or zoonotic transfer. For combating these infections immunological defense systems trigger phagocytosis or apoptosis, produce cytokines or antibodies, and release inflammatory or cytotoxic mediators (Andersen et al., 2021). Nonetheless, the impact of infectious diseases is still substantial in low and lower-middle income countries, and the mortality and morbidity attributable to neglected tropical diseases (NTDs) is also high (Baker et al., 2022) of which Leishmaniasis is believed to have a high mortality rate in humans and is further correlated to the disadvantages of current medications, such as their high toxicity and drug resistance (J et al., 2021). Leishmaniasis is an intracellular protozoal infection that is chronic and prolonged and is caused on by multiple species of the genus *Leishmania*. Evidences point towards leishmaniasis could become more common due to factors like climate change, the discovery of effective vectors and reservoirs, a highly mobile population, significant population groups with documented exposure histories, HIV, and the widespread use of immunosuppressive drugs and organ transplants encouraging potential for sustained autochthonous spread (Curtin and Aronson, 2021). Apart from standard drugs and FDA-approved drugs such as pentavalent antimonials, meglumine antimoniate, sodium stibogluconate, Pentamidine, Liposomal amphotericin B and Miltefosine, (J et al., 2021) novel therapeutic approaches are also progressing and gaining a lot of attention such as phytotherapy, nanotherapeutics, metabolite encrichment, Antimicrobial peptides (AMPs), Proteasome and epigenetic modifiers (Sundar and Singh, 2018).

The chromatin remodeling in the parasite is thought to be significantly influenced by altered epigenetic histone modifications. It’s intriguing to note that the parasite alters host gene expression as well, enabling the host immune response to be suppressed or hijacked. The silencing of genes specific to macrophages that are involved in defense against these parasites has been linked to epigenetic factors such DNA methylation of cytosine residues (Afrin et al., 2019). Many studies highlight the potential of epigenetic factors as a prime target for vaccine as well as therapeutic target development. Recombinant histone H1 vaccination of monkeys resulted in a slower development of cutaneous lesions compared to controls suggesting histone H1 can be a candidate for vaccine development against cutaneous leishmaniasis (CL) in humans (Masina et al., 2003). It was also reported that, rLdH2-4 exclusively generates Thl-type immune responses, providing significant protection against experimental visceral leishmaniasis that cured patients/endemic contacts, and cured hamsters (Baharia et al., 2021). It was also shown that, cutaneous lesions exhibit amplified matrix metalloproteinase 1 (MMP1) which may be regulated epigenetically by Friend leukemia virus integration 1(FLI1) and Interleukin 6 (IL-6) suggesting FLI1 can be a therapeutic target (Almeida et al., 2017). Promising therapeutic targets might include the enzymes responsible for histone post translational modifications, particularly those that contain epigenetic reader modules and bromodomains such as Sirtuins of *L.donovani* (Afrin et al., 2019). The effect of imipramine on IL-10/IL-12 axis reciprocity was studied for clearance of antimony resistant *L.donovani* which specifically targeted HDAC11 which inhibited its ability to acetylate IL-10 promoter region leading to downregulation of IL-10 (Mukherjee et al., 2014). Even though these vaccines and drugs with potentials to inhibit parasite growth have been reported, none of them has been introduced in the market specifically targeted towards any form of leishmaniasis. The short comings and limitations of these molecules is the phenotypic response of immune cells. In order to understand drug designing it is important to understand immune cell population as a network and dissecting the time dependent response of the host immune cell towards parasite. Hence population dynamics, cell enrichment, phenotype characterization and heterogeneity distribution may lay an insight as to which subset of cells are playing role in parasite proliferation and which genes are supporting the parasite survival process.

Majority of single-cell profiling investigations have so far relied on quantifying RNA by sequencing (scRNA-seq). While this offers glimpses of the inter- and intracellular variation in gene expression, investigation into the epigenomic landscape in single cells has enormous promise for revealing a key component of the regulatory logic of gene expression programmes (Chen et al., 2019). Together with help of recent developments in array-based technologies, droplet microfluidics, and combinatorial indexing through split-pooling, the single-cell Assay for Transposase Accessible Chromatin using sequencing (scATAC-seq) has been able to generate chromatin accessibility data for thousands of single cells in a relatively simple and affordable way (Chen et al., 2019). Conventional methods that use bulk tissue samples as input lack the resolution to assess the temporal dynamics of cell-type-specific use. This methodology has been successfully implemented to embryonic tissues in *D. melanogaster*, developing mouse forebrains, adult mouse tissues, (Fang et al., 2021) human pancreatic islets (Rai et al., 2020), human testicular cells (Wu et al., 2022), fetal human retina (Mulqueen et al., 2021) and mouse cardiac progenitor cells (Jia et al., 2018). This technique enables to separate cells by discriminating them on the basis of cell types; source and cell variability to generate population clusters (Pott and Lieb, 2015).

scATAC-seq is based on bulk ATAC-seq which involves isolation of nuclei from single cells in the sample through FACS. Hyperactive Tn5 transposase catalyses the process of tagmentation by integrating sequencing adaptors to the targeted DNA by initiating its binding to the DNA followed by release from DNA post tagging through heat or denaturing molecules. While nuclei are still intact, single nuclei are isolated (Xu et al., 2021). The nuclei are later lysed and loaded onto 96-well plates containing specially barcoded transposases, then sorted again before being dispensed into a second 96-well plate for fluorescence-activated cell sorter (FACS). Later, second set of barcodes known as unique molecular identifiers (UMIs) are introduced in the amplification step (Zhu et al., 2020). By identifying distinct combination of both the barcode combinations, around 1500 cells with a median range of 2500 and 11% collision rate can be read (Baek and Lee, 2020). The downstream analysis may result in cell clustering that distinguishes between distinct cell types in a mixed cell population and find peaks that are more or less accessible to particular cell types (Baker et al., 2019) thus identifying complicated cell populations, connecting regulatory elements to their target genes, and mapping regulatory dynamics during complex cellular differentiation processes through the chromatin-regulatory landscape on the cell clusters (Rai et al., 2020).

Epigenetic modifications and chromatin remodeling framework of macrophages with respect to *L.major* infection that causes CL is still poorly understood. Macrophage being the most prime target of infection may dictate disease fate and the modular characteristics of parasite may influence the host macrophage plasticity for survival (Khandibharad et al., 2022). Fine balance between different types of macrophages such as M1 which mediates Th1-type response and M2 macrophages having different subsets such as M2a, M2b, M2c and M2d that elevates Th2-type response regulates the parasite clearance (Tomiotto-Pellissier et al., 2018). One of the key mechanisms of parasite survival which we had reported was through governing IL-10 and IL-12 reciprocity. Through computational and systems biology integrated framework models we had previously reported that epigenetic factors such as NFAT5 and their regulators mainly SHP-1 may play deterministic role in regulating IL-10 and IL-12 reciprocity (Khandibharad and Singh, 2022a). In this work we unravel the epigenetic paradigm and chromatin remodelling framework through scATAC-seq of *L.major* infected RAW264.7 mouse macrophage cell line at different time scales. We have identified motifs which dictate the reciprocal regulation of cytokines, parasite survival response and infectivity. We assume that the time lapse shift in population dynamics, cluster heterogeneity and change in peak of motifs are occurring due to change in dynamicity of chromatin accessibility. We are also for the first time reporting the quantified data of macrophage infected cells at single cell level and we are coining the term “sleepy macrophages” which are not even expressing housekeeping genes but are exclusively expressing NFAT5. We also suspect that precision targeted therapeutics that can modulate phenotype population and might help in resolving disease.

## Methodology

### Sample preparation

The ability of macrophages to phagocytose and promote parasite growth makes them prime resident cells for *Leishmania* although these cells do function as the primary effector cells in the clearance of infection (Liu and Uzonna, 2012). As a component and regulator of adaptive immunity, macrophage facilitate expression of interferon-gamma (IFN-γ) and tumor necrosis factor-alpha (TNF-α), interleukin 12 (IL-12) that facilitates production of Nitric oxide (NO). To endure in the hostile macrophage environment, *Leishmania* subverts the macrophage cellular process, metabolic pathways molecular functions and promotes chromatin remodelling to produce immunosuppressive molecules such as transforming growth factor β (TGF-β), interleukin 10 (IL-10) for its survival (Loría-Cervera and Andrade-Narvaez, 2020).

*L.major* exhibits unique epigenetic modifications that are not reproduced by any other *Leishmania spp*. (Kamhawi and Serafim, 2020). Our previous findings had reported reciprocal expression of two crucial cytokines IL-10 and IL-12 and we identified the time points of their distinct expression (Khandibharad and Singh, 2022a) (Khandibharad and Singh, 2022b). Therefore we submitted four samples for study. Mouse derived stable macrophage cell line RAW264.7 cells, 1X10^6^ were infected with stationary phase *L.major* promastigotes, and were incubated for 6hr, 12hr and 18hr. Post incubation, cells were scrapped off and cryo preserved in cryomix [90% Fetal Bovine Serum (FBS) + 10% Dimethyl sulfoxide (DMSO)], control used for study was uninfected RAW264.7 cells. To ensure maximum revival capacity of cells, they were frozen with gradual temperature changes; 0°C for 30 minutes, −20 °C for 3hrs, −80 °C overnight and liquid nitrogen storage for 24hrs. Each sample was submitted to Neuberg Centre for Genomic Medicine, Supratech Reference Laboratory in duplicates for nuclei isolation, microfluidics based library preparation and sequencing.

### Sample Quality check

The cryo vials were revived by thawing them at 37 °C in water bath for 2 mins, followed by mixing with 10ml pre-warmed media and centrifuged at 300rcf for 5min. The cell pellet was resuspended in 1XPBS+ 0.04% BSA (Sigma #A2153) and passed through 40μm Flowmi Cell Strainer (Sigma NB.01). To check the viability of cells post revival, gently 20 μL of the sample and 20 μL of 0.4% trypan blue (ThermoFisher #15250061) were mixed, 10 μL of the mix was loaded into CountessTM cell counting chamber slides (Thermofisher #A51876). Cells were counted using Countess™ 3 FL Automated Cell Counter (ThermoFisher).

### Nuclei Isolation and Quality check

Utilizing the CG000169 procedure from 10X Genomics, nuclei isolation was performed. The revived cells were centrifuged at 300 rcf for 5min at 4 °C. To that 100 μL of chilled Lysis buffer [10mM Tris-HCl (pH 7.4), 10mM NaCl, 3mM MgCl_2_, 0.1% Tween-20, 0.1% Nonidet P40 Substitute, 0.01% Digitonin and 1% BSA] was added and gently mixed 10 times and reaction mix was incubated on mice for 5 minutes. To the lysed cells, 1ml wash buffer [10mM Tris-HCl (pH 7.4), 10mM NaCl, 3mM MgCl_2_, 1% BSA and 0.1% Tween-20] was added and mixed gently 5 times followed by centrifugation at 500 rcf for 5 min at 4°C. Supernatant was discarded and nuclei pellet was resuspended in 7 μL of chilled Nuclei buffer. Fifth portion of 10 μL nuclei suspension buffer was made with nuclei suspension and nuclei buffer and was mixed with 10 μL of 0.4% trypan blue stain of which 10 μL of the mix was loaded on to the slide chamber. Nuclei concentration and viability was quantified using Countess™ 3 FL Automated Cell Counter.

### Transposition of isolated Nuclei

Nuclei suspensions were incubated with Transposase that was present in Transposition mix. Transposase fragments the DNA at open chromatin regions by entering the nuclei. Adapter sequences are simultaneously linked to the ends of the DNA fragments through Polymerase chain reaction (PCR). The reaction mix was incubated at 50 °C for 30min, 37 °C for 30min and 4 °C for hold.

### Gel beads-in-emulsion and barcoding

Initiation of the steps was done by using CG000496 protocol. In brief, Barcoded gel beads, transposed nuclei, a master mix and portioning oil were loaded on a chromium Next GEM Chip H are combined to achieve single cell resolution using Chromium iX. The distribution of the nuclei occurs at a limiting dilution such that most (90-99%) of resulted GEMs have one or no nuclei, while the majority of the rest do. The Gel Beads dissolves following GEM synthesis. After mixing and released from (1) an illumina P5 sequence, (2) a 16 nucleotide 10X Barcodes and (3) a Read 1 (Read 1N) oligonucleotides are further taken for thermal cycling to generate 10X barcoded single stranded DNA. The samples for thermal cycler based extension were incubated at 72 °C for 5mins, 98 °C for 30s, 98°C for 10s, 59°C for 30s, 72 °C for 1min, this cycle takes place with 12 times repetition and finally is kept at 15°C to hold.

### Single cell library preparation and sequencing

During library preparation P7 and a sample index are added through PCR. The P5 and P7 sequences used in Illumina^®^ bridge amplification were present in the final libraries. Following the manufacturer’s instructions, final libraries were quantified using a Qubit 4.0 fluorometer (Thermofisher #Q33238) and a DNA HS test kit (Thermofisher #Q32851). We scanned the library on the Tapestation 4150 (Agilent) using high sensitive D1000 screentapes to determine the insert size. Final QC checked libraries were sequenced on Illumina (Novaseq 6000) using SP flowcell at 50:8:16:50 cycles. Post Sequencing the data is demultiplexed using the cell ranger arc V7.0 and BCL2FASTQs were futher processed to generate RAW FASTQ, websummary files and clope file using mm10 reference genome.

### Clustering

Using log2 value as a filter parameter, custom filters were made to identify the population heterogeneity. We used loupe browser for cell clustering. Peak calling is frequently repeated for each cluster in order to determine the accessible chromatin regions for various cellular populations. These regions are then the subject of a statistical test for correlations with different pre-defined genetic features. The primary objectives of downstream analysis techniques are to identify novel regulatory components and comprehend how they function within a cell (Baek and Lee, 2020).

### Dimensionality reduction for identification of principle components (PC)

A key technique for examining scATAC-seq datasets is through principle component analysis (PCA). Reduction of data dimensions helps in big data processing, interpretation and identifying uniqueness of each component. We used Factoextra and FactoMineR to perform PCA. We considered the data table as X and transformed it to the original coordinate system by orthogonal linear transformation. Let Fs (or Gs) stand for the vector representing the coordinates for the rows (or columns) on the axis of rank “s”. According to the so-called “transition formulas,” these two vectors are connected and represented as

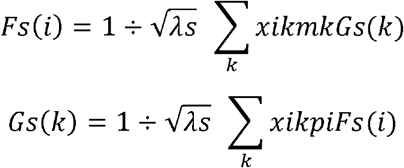

where Fs(i) is the coordinate of individual i on axis s, Gs(k) is the coordinate of variable k on axis s, λs is the eigenvalue associated with axis s, mk is the weight assigned to variable k, pi is the weight assigned to individual i and xik is the catchall concept of the data table (row i, column k) (Lê et al., 2008). As clusters 3 and 6 from 6hr infected sample were identified as unique clusters, we identified the principle components from 6hr sample.

### Gene Set Enrichment Analysis (GSEA)

GSEA enables evaluation of gene expression data at levels of gene sets resolving the concerns arising due to insignificant statistics of gene expression post analysis, non-unique biological activity, identifying single gene function in cellular process and prevents overlaps among similar kind of studies (Subramanian et al., 2005). A gene set, or a list of genes known to be involved with a specific biological activity, gene ontology (GO), molecular function, or pathway, is compared to the ranked gene list. The metric is required to determine the enrichment score (ES), which measures how much a gene set is overrepresented at the extremes of the ranking list (Zito et al., 2021). We took into account the GSEA’s capability to evaluate 814 genes in order to provide physiologically pertinent details regarding the differential expression of genes belonging to cluster 3. By classifying the clusters into two groups, Diseased and Normal (1, 0 respectively), we investigated the differential expression and enrichment of genes from 6hr infected sample in cluster 3 and 6 and compared to clusters 1, 2, 4 and 5in the dataset.

### Differential correlation analysis

It’s critical to understand how the correlation between molecules under two different experimental conditions has changed, in addition to how the mean amounts of molecules in the omics data have changed. We employed DiffCorr package, to understand the correlation among the clusters in the samples. For each dataset, DiffCorr generates correlation matrices, locates the first principal component-based “eigen-molecules” in the correlation networks, and uses Fisher’s z-test to assess differential correlation between the two groups (Fukushima, 2013). We performed Differential Correlation analysis for 6hr sample.

### Construction of Transcription Factor Target Gene (TFTG) Network

As the transcription factors which are expressed in Cluster 3 will result in upregulating the genes from the same cluster, we prepared an integrated framework network of the gene targets and transcription factors enriched in cluster 3. We used the TF link database, which offers thorough and extremely reliable information on transcription factor - target gene interactions for numerous organisms, including *Mus musculus*, for this purpose. It integrates information from other TF databases, including JASPAR and TRRUST database, to offer cumulative statistics for the TFs for a specific organism.

The entire integrated biomolecular interaction network of Transcription factor-Target gene (TFTG) was constructed and analysed using Cytoscape (v. 3.6.0). The initial TFTG network had 814 genes obtained from cluster3 and 21 transcription factors. The constructed TFTG network consisted of 476 nodes and 1060 edges. This initial network was then subjected to simulated annealing algorithm in Cytoscape which will give a robust inter-regulatory TFTG network that is resilient and comprehensible in nature; as the loosely connected edges of the network are filtered out. The most clustered or heavily weighted nodes are positioned at the bottom of the network using the simulated annealing process, which analyses each node in the network. In contrast, nodes with lesser clusters are arranged in descending order in the upper part of the network.

The robust simulated annealing network obtained is further analysed considering the potential of Cytoscape plugin - CytoHubba. CytoHubba uses a double screening scheme for ranking nodes and edges in a network. Further network analysis through CytoHubba helps us understand the function of an individual node and its collaboration with other nodes in a cluster. It uses algorithms like Betweenness Centrality, Closeness Centrality, Degree of Nodes, Maximal Clique Centrality (MCC), Clustering Coefficient, BottleNeck and others to present a condensed and more robust nature of the inter-regulatory transcription factor network with their putative target genes.

## Results

### Single cell ATAC sequencing reveals macrophage clusters post *L.major* infection

After analysis of all the samples, 6 cell clusters were observed in control and 6hr sample, 12 clusters in 12hr sample and 10 clusters in 18hr samples (Supplementary file 1 (S1, S2 and S3). These clusters had different cell number with varied percentage population. There was increase in the number of cell clusters from 6hr to 12 hr although the number of cluster decreased at 18hr as compared to 12hr sample.

### Percentage population of M1 and M2 macrophages changes with time point of infection

M1 and M2 subtypes macrophages were identified based on their differential expression levels for which threshold for marker genes were set with cut off log2>0.5. For M1 we used IFNγ and CD80 as markers as they characterize high pro-inflammatory response. M2 subtypes commonly express IL10 hence to get an overview of entire M2 we used IL10 as a marker for which threshold was set with cut off log2>0.5 (Figure 1, 1a-1d). M2 subtypes which include M2a, M2b, M2c and M2d were characterized by markers. Parameters for M2a were set as log2 CCl24>1 and log2 SOCS3>1, for M2b log2 TNF>1 and log2 CCl1>1 for M2c, log2 TGFB>1 and log2CD163>1 and for M2d log2 NOS2>1 and log2 CCl5>1 (Khandibharad et al., 2022) (Figure 1 (2)). We could see the shift in macrophage population with time points of infection where M2 and M2a population was most dominant in all the samples, M1 was most dominant in 6hr infected sample, M2d was enriched in 6hr infected sample as well. In 12hr infection sample a decline in number of cells was observed for all the macrophage types and subtypes with a gradual increase in 18hr (Figure 1 (3)).

**Figure 1:**
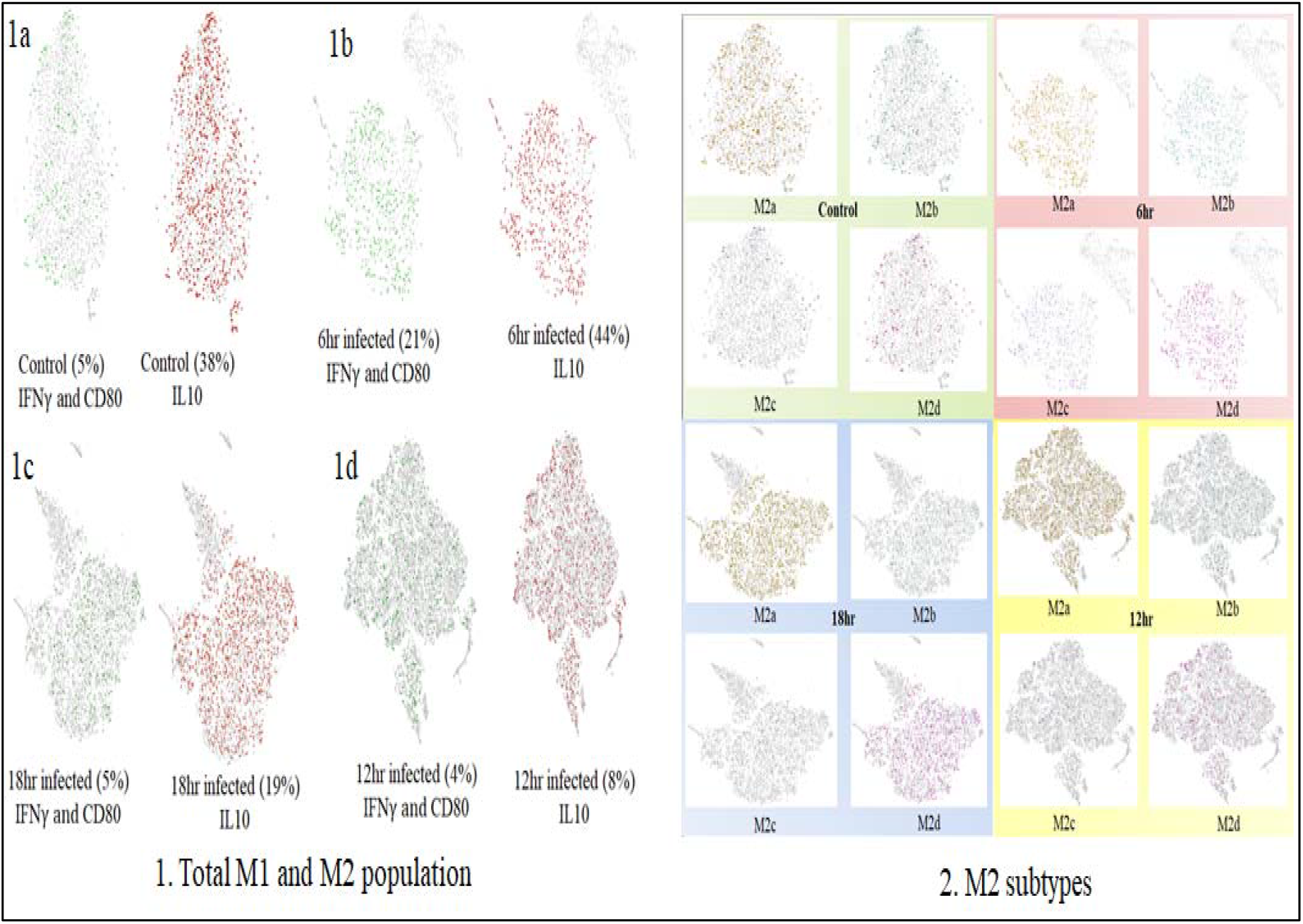
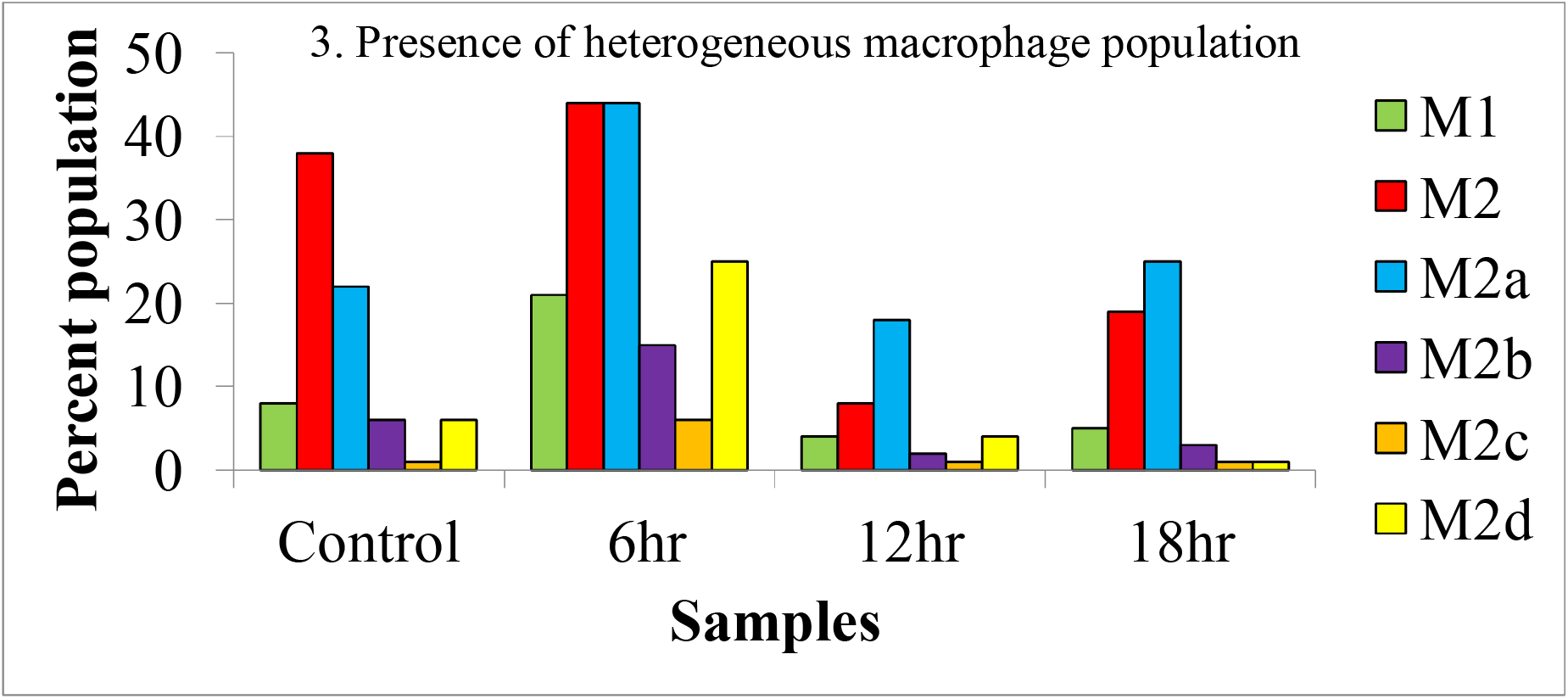
(1) Expression of M1 and M2 markers IFNγ, CD80 and IL10 in samples. 1a-1d Control, 6hr, 12 hr and 18hr. (2) Percentage of M2 subtype population. (3) Graphical representation of overall macrophage types and subtypes in all the samples.

### Infection of macrophages with *L.major* differentiates IL10 and IL12 production population

IL12 being pro-inflammatory cytokine and IL10 anti-inflammatory are explicitly expressed as response to *L.major* infection by macrophages (Figure 2 (2)). Populations were identified based on unique expression of IL12 more than IL10 and vice versa. This categorization identified the cellular response to infection with time. The parameter used was log2IL12b>0.5 and log2 IL10<0.5 for distinguishing IL12 producing group from IL10 producing group for which parameter for filtering was set as log2 IL10>0.5 and log2 IL12b<0.5 (Figure 2 (1a-1d)). In the uninfected sample we observed that population expressing IL10 were more than IL12 expressing cells although we observed slight changes in depletion of IL10 producing cells at 6hr of infection. At12 hr, Il12 expressing cells increased but did not dominate over IL10 expressing cells and they diminished at 18hr. We could infer that, may be at 6hr and 12hr of infection macrophages tries to supress IL10 production and favour IL12 production but due to modulatory effects of intracellular *L.major* on regulation of host cytokine machinery, macrophages fail to produce IL12 over IL10 and therefore reduces eventually.

**Figure 2:**
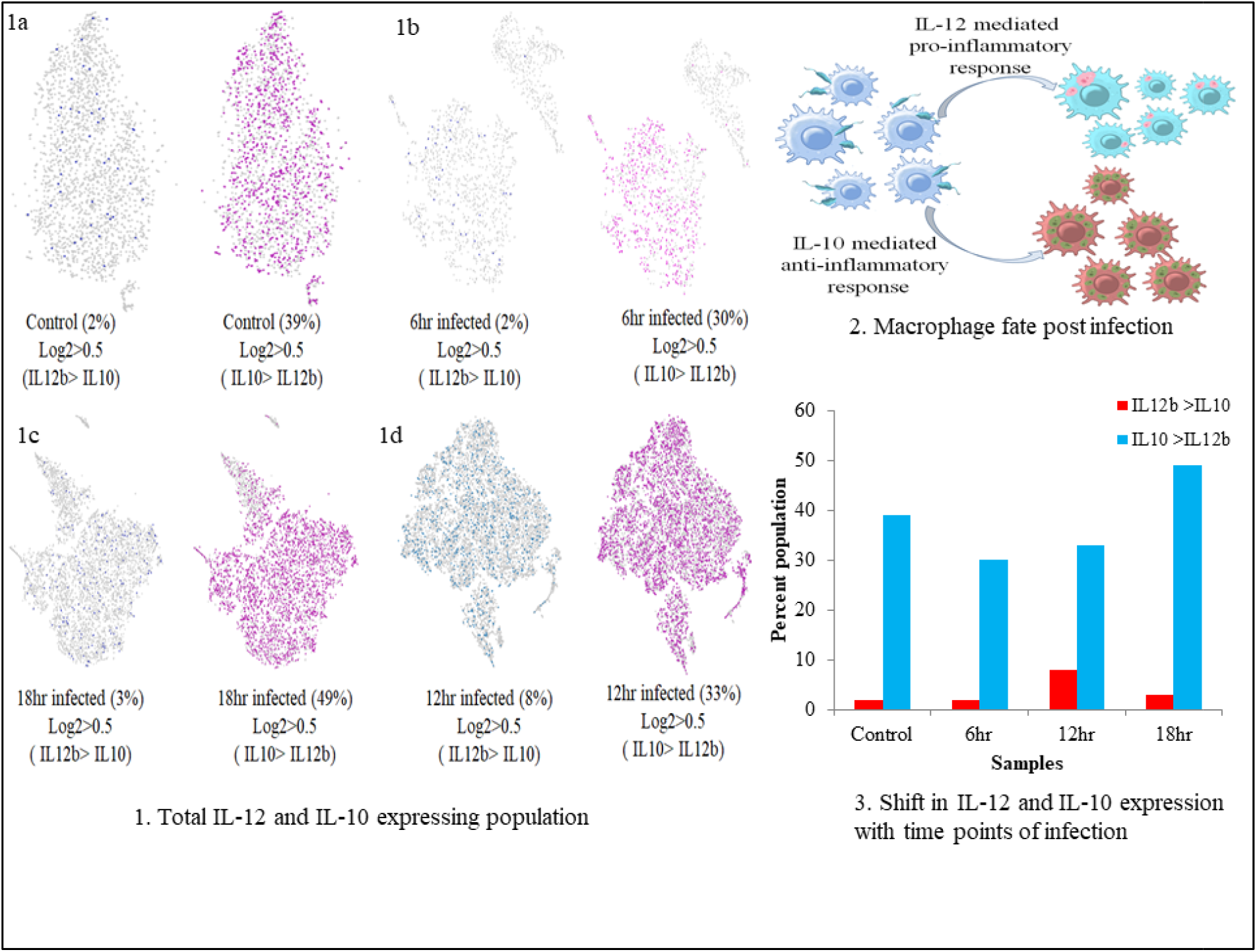
Identification of reciprocal relationship between IL10 and IL12. (1) Macrophage expressing IL12 more than IL10 and vice versa in all the samples (1a-1d) Control, 6hr, 12hr and 18hr. (2) Abstract highlighting the fate of macrophages post infection with *L.major* if IL12 and IL10 are secreted. (3) Change in expression patterns of IL10 and IL12 with time.

### Reciprocal regulation of IL10 and IL12 through NFAT5 and SHP-1 expression

From our previous findings we have already reported that upon *L.major* infection NFAT5 can govern expression of IL12 and nitric oxide and SHP-1 can regulate NFAT5 through dephosphorylating it at auxiliary export domain which inhibits NFAT5 dependent pro-inflammatory response and directs the cellular machinery towards IL10 and Arginase mediated ornithine cycle (Figure 3 (3)). We identified if these deterministic genes are expressed together to get an insight into IL10 and IL12 expression axis. Parameters used to identify these populations across samples were set as log2 NFAT5 motif>0.5, log2 IL12b sum>0.5 and log2 NOS>0.5. For second group the distinguishing parameter were set as log2 IL10>0.5, log2 ptpn6>0.5 and log2 Arg1>0.5 (Figure 3 (1a-1d)). We could discern that at 6hr of infection parasite eliminating population was dominating over parasite survival favouring population. At 12hr population architecture dropped without changing the dynamics however, at 18hr the dynamics changed with subtle shift of parasite survival favouring population over parasite eliminating population (Figure 3 (2)).

**Figure 3:**
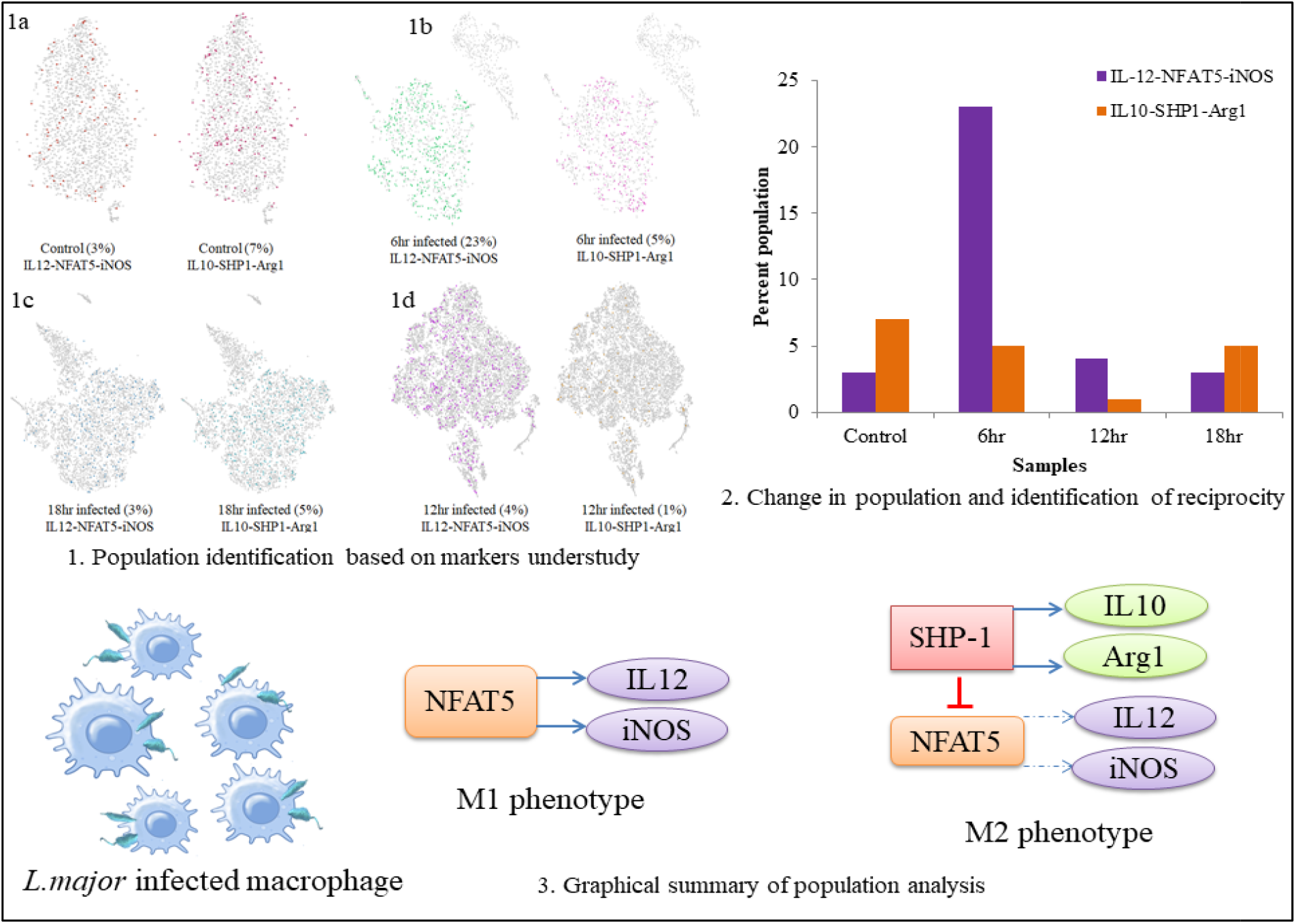
Identification of reciprocity in IL10 and IL12 expression patterns through NFAT5 and SHP-1. (1) Populations expressing IL12, NFAT5 and iNOS versus populations expressing IL10, SHP-1 and Arginase1 (1a-1d) Control, 6hr, 12hr and 18hr. (2) Graphical representation highlighting reciprocal relationship between IL10 and IL12 mediated by NFAT5 and SHP-1. (3) Pictorial abstract of the aim behind the analysis.

### Infection with *L.major* for 6hr identifies sleepy macrophages

As we observed the changes in gene expression pattern which shifted the cell population dynamics from parasite eliminating group to parasite survival cell subset starting at 6hr post infection, we analysed the chromatin accessibility of IL10, IL12p40, SHP-1 and NFAT5 and used B-Actin and GAPDH as housekeeping gene control. To our surprise, there were two clusters which showed minimal expression of housekeeping genes and genes under study (Figure 4 (A-E)) (Table 1). This made us intrigued as no other clusters from other samples showed similar expression pattern. As these cells are alive, the chance that these undergo a survival mechanism in order to combat the infection was fascinating. NFAT5 motif which regulates IL12 and IL10 expression was highly enriched (Figure 4 (F)) suggesting its involvement through molecular mechanisms, cellular processes and biological signaling may play a role in these cells to govern IL10 and IL12 reciprocity. Results for all samples is in (Supplementary file 1 (S4))

**Figure 4:**
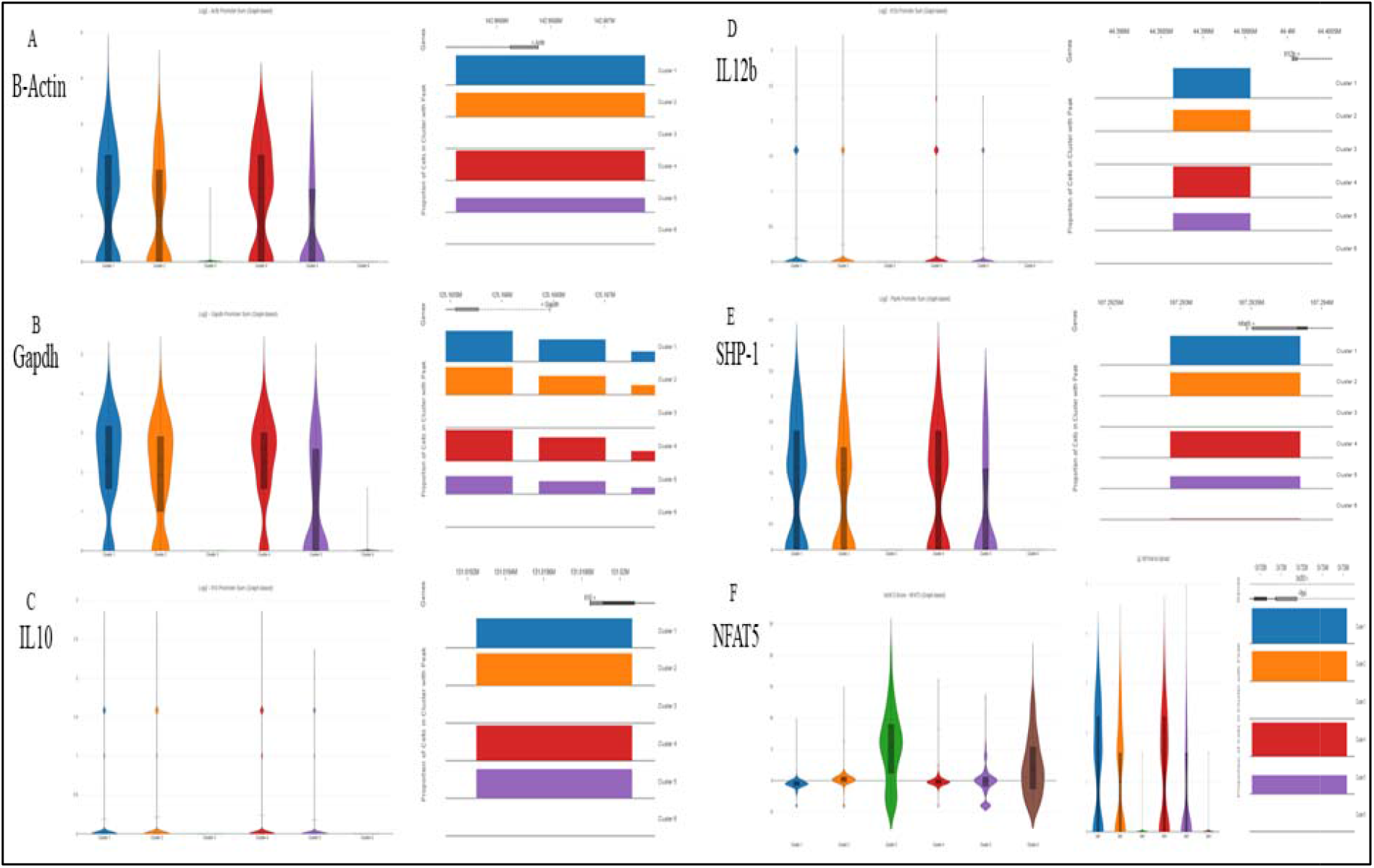
Differential expression of genes in 6hr sample in all clusters. (A-E) Violin plot and peaks of β-Actin, GAPDH, IL10, IL12 and SHP-1. (F) Violin plot of NFAT5 motif accessibility, Violin plot and peak of NFAT5.

**Table 1:**
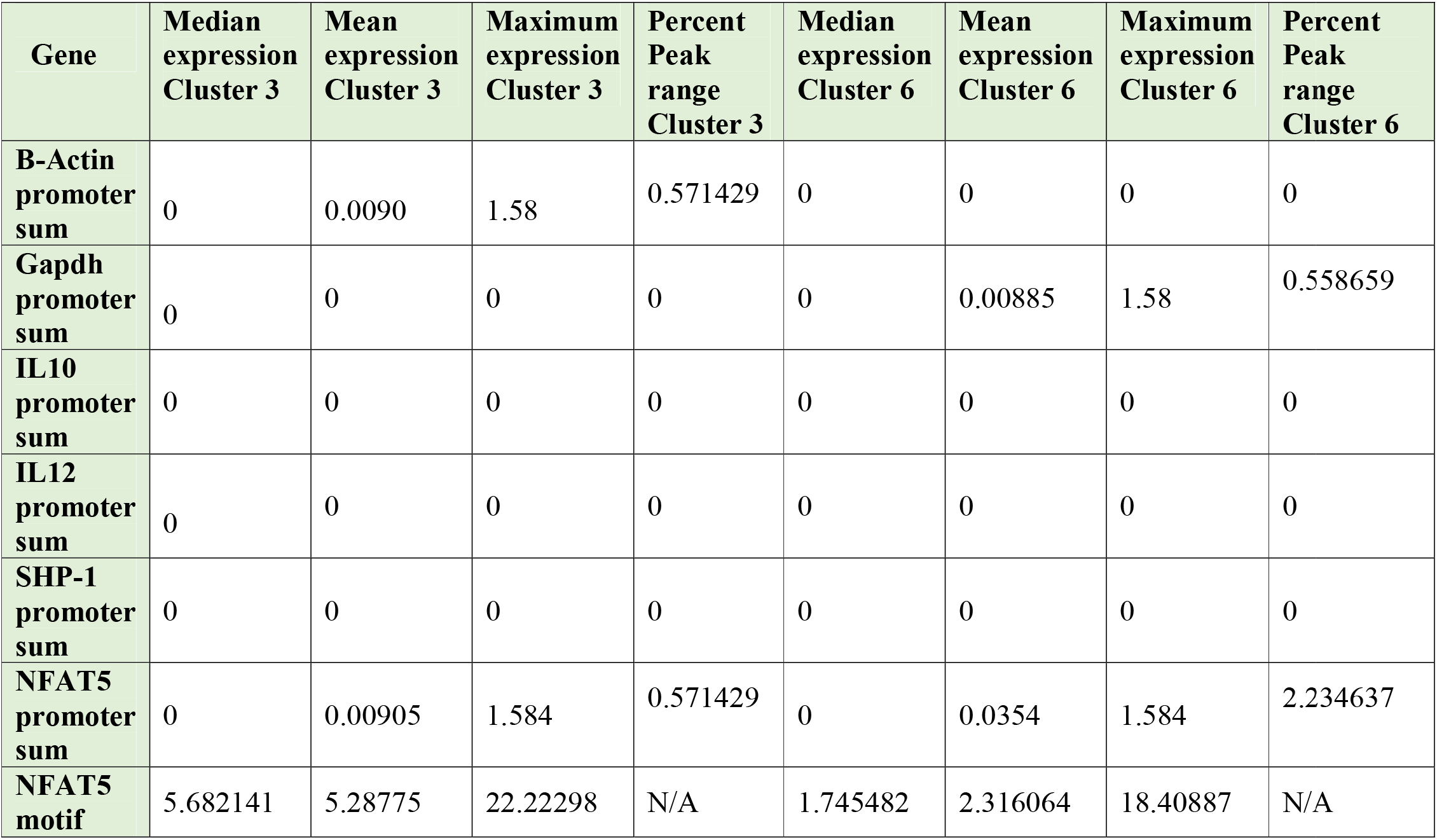
Expression and accessibility of gene promoters under study.

| Gene | Median expression Cluster 3 | Mean expression Cluster 3 | Maximum expression Cluster 3 | Percent Peak range Cluster 3 | Median expression Cluster 6 | Mean expression Cluster 6 | Maximum expression Cluster 6 | Percent Peak range Cluster 6 |
| --- | --- | --- | --- | --- | --- | --- | --- | --- |
| B-Actin promoter sum | 0 | 0.0090 | 1.58 | 0.571429 | 0 | 0 | 0 | 0 |
| Gapdh promoter sum | 0 | 0 | 0 | 0 | 0 | 0.00885 | 1.58 | 0.558659 |
| IL10 promoter sum | 0 | 0 | 0 | 0 | 0 | 0 | 0 | 0 |
| IL12 promoter sum | 0 | 0 | 0 | 0 | 0 | 0 | 0 | 0 |
| SHP-1 promoter sum | 0 | 0 | 0 | 0 | 0 | 0 | 0 | 0 |
| NFAT5 promoter sum | 0 | 0.00905 | 1.584 | 0.571429 | 0 | 0.0354 | 1.584 | 2.234637 |
| NFAT5 motif | 5.682141 | 5.28775 | 22.22298 | N/A | 1.745482 | 2.316064 | 18.40887 | N/A |

### Identification of principle components from sleepy macrophages

To identify which genes are enriched in Cluster 3 and Clsuter 6 we extracted entire feature from 6hr sample and observed the genes which are upregulated in cluster 3 and Clsuter 6 are significantly downregulated in Cluster 1,2,4 and 5 (Supplementary file 2 (S5)). From the scree plot we observed the elbow forming Cluster 3 there after the slope was flat suggesting cluster 3 has more variance and can be taken further to identify principle components (Figure 5 (A)). Cluster3 had higher cos2 when indiviual genes were analyzed although number of variables of the same were less (Figure 5 (C-D)). Higher the cos2 value more important is the individual feature as a PC, based on level of significance we identified PCs (Figure 5 (B and E)) (Supplementary file 2 (S7)).

**Figure 5:**
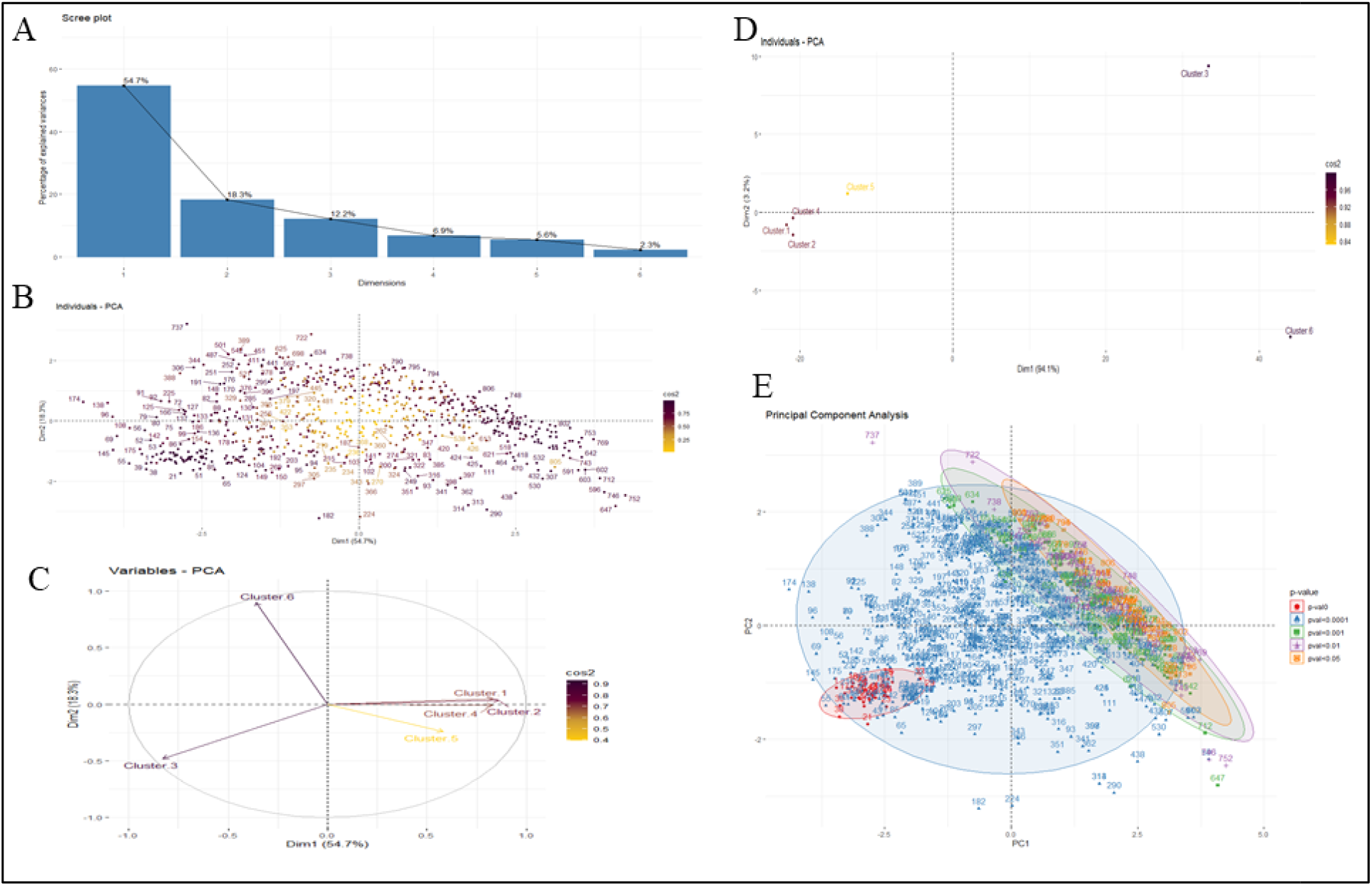
Principle component analysis of 6hr sample by differential expression of all the clusters. (A) Scree plot, (B) Genes PCA, (C) Variables in clusters, (D) PCA of clusters and (E) PCA based on p-value.

### Gene set enrichment and differential correlation of sleepy macrophages

The main outcome of the gene set enrichment analysis is the enrichment plot illustrating the enrichment score (ES), which measures the degree to which a gene set is over-represented at the top or bottom of an ordered (ranked) list of genes. In the enrichment plot the magnitude of the increment in the enrichment plot depends on how well the gene correlates with the defined phenotype (class), the graph here illustrates the positive enrichment of gene sets showing rapid increase in the expression of genes belonging to cluster 3 (the diseased class) with an enrichment score of greater than 0.8. Additionally, it has been observed that the leading-edge subset of a gene set (subset of members contributing most to the ES) positively correlates with the disease phenotype. Furthermore, the bottom-most portion of the enrichment plot shows the value of the ranking metric which measures every gene’s correlation in the gene set with the defined phenotype. The ranking metrics’ value goes from positive to negative as we move down the ranked list; positive value indicating correlation with the first phenotype (diseased) and a negative value indicating correlation with the second phenotype (normal). With respect to the ES the highest-ranking genes from the gene set fall between ranks 0 - 100 ranks (Figure 6 (B)).

**Figure 6:**
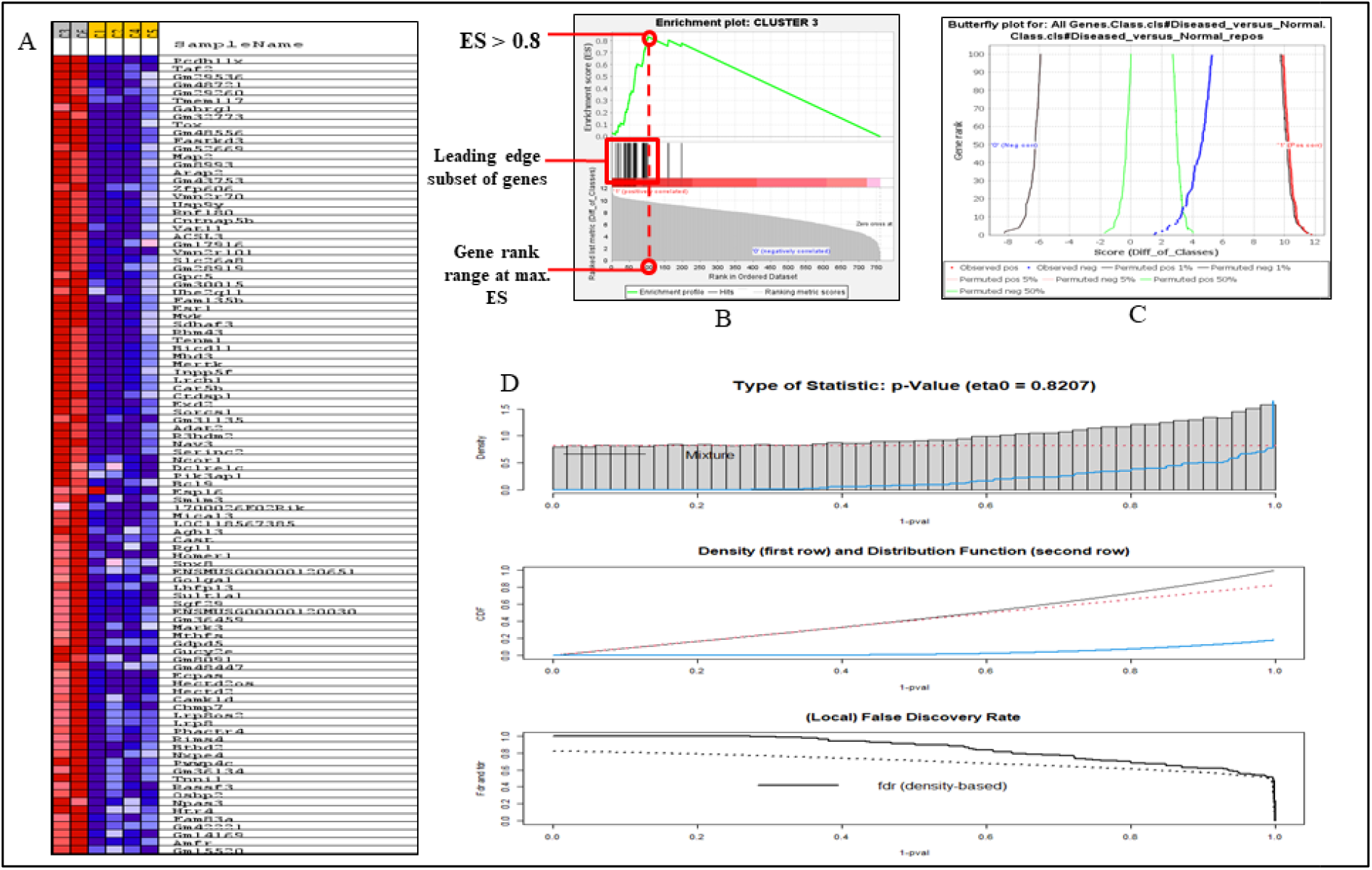
Gene enrichment analysis from Cluster 3 and Cluster 6 of 6hr sample. (A) Heat map of genes enriched from cluster 3 and 6. (B) Enrichment plot of Cluster 3 (C) Butterfly plot of Cluster 3 and (D) Differential correlation of Cluster 3 with other clusters.

Alongside, the butterfly plot shows the positive and negative association of genes for each gene in the ranked list according to their rank and ranking metric score. It shows the observed correlation for the top genes as well as permuted (1%, 5%, 50%) positive and negative correlation. The plot depicts the observed negative genes belonging to the normal phenotype to be negatively co-related via aberrant expression shifting more towards the disease phenotype supporting the enrichment plots (Figure 6 (C)). Moreover, a heat map is also used to depict the (clustered) genes in the leading-edge subgroups. The correlation between the ranked genes and the designated class is displayed on a heat map for each phenotype. Expression values are depicted on the map as colors; with the hues (red, pink, light blue, dark blue) according to the expression value range (high, moderate, low, and lowest). The heat map in this case demonstrates that cluster 3 and 6 have higher expression of the leading-edge subset of genes correlating the diseased phenotype as compared to other clusters in the dataset (Figure 6 (A)). To observe the gene differential expression pattern change between Clusters 1,2,4,5 and Cluster 3&6 we performed (Supplementary file 2 (S11)) differential correlation of differential gene expression patterns using Pearson’s correlation coefficient and observed density (p-value) and distribution function to be correlated, with false discover rate to be decreasing with p-value (Figure 6 (D)).

### TFTG network links cellular pathways and novel markers in sleepy macrophages

Our comprehensive study identified 51 transcription factors enriched in sleepy macrophages which may regulate 814 identified genes, we could map 22 transcription factors and their target genes from sleepy macrophages genes and constructed the network which had 476 nodes and 1060 edges (Figure 8 (B and C) (Supplementary file 2 (S10)). The initial network was subjected to simulated annealing algorithm to improve the network’s robustness and comprehensibility making the network distribution lucid for understanding which further depicted the reduction in the number of multi-edge node pairs which indicates how frequently the adjacent/neighbouring nodes are connected by more than one edge; the simulated annealing algorithm filtered out weakly connected edges making the network more robust (Table 2) and distinctly highlighting the densely connected nodes/transcription factors namely Mef2c, Isl1, Rxra, Jun, Pparg, Ascl1 and Onecut2 (Figure 8 E).

**Figure 7:**
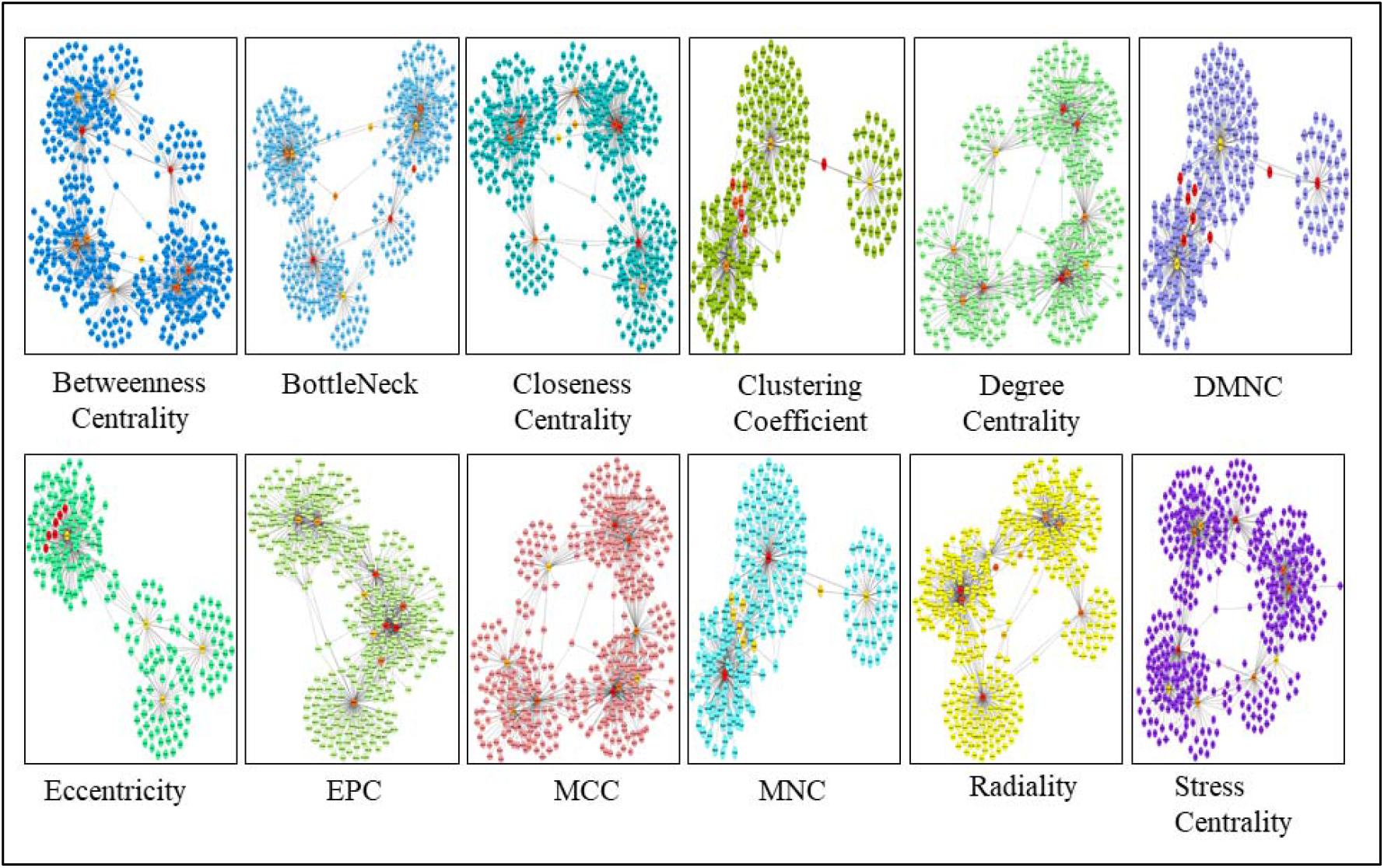
Substantial modules based on 12 scoring techniques were identified from the leading inter-regulatory TFTG network.

**Figure 8:**
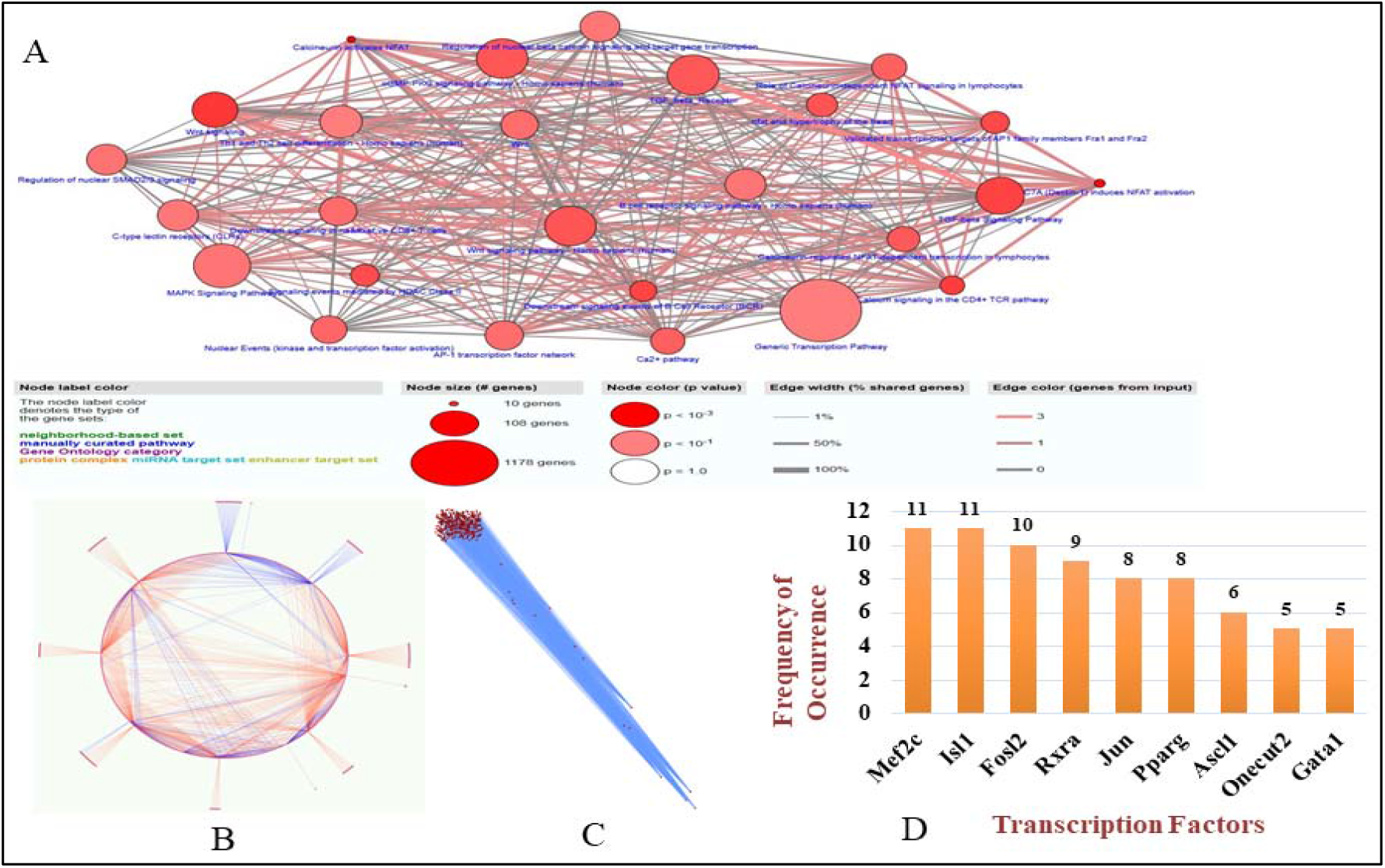
TFTG network analysis. (A) Pathway enrichment of sleepy macrophages genes and transcription factors (B) The simulated network’s circular layout demonstrating the strength of chosen Transcription Factors over the whole network. (C) The inter-regulatory transcription factor - target genes (TFTG) network after running the simulated annealing algorithm, showing placement of heavily weighted nodes (TFs) positioned at the bottom of the network and (D) The top 5 ranked Transcription factors are represented graphically, based on their frequency of occurrence according to the 12 scoring techniques of CytoHubba plugin.

**Table 2:** Statistical Analysis of Transcription Factor - Target Genes (TFTG) inter-regulatory network after Simulated Annealing Representing a Decrease in the Multi-edge node pairs, making the Network More Robust by filtering out the loosely connected edges.

| Parameters | Original Network Values | Simulated Annealing Network Values |
| --- | --- | --- |
| Clustering Coefficient | 0.005 | 0.005 |
| Connected Components | 1 | 1 |
| Network Diameter | 8 | 8 |
| Network Radius | 4 | 4 |
| Shortest Paths | 226100 (100%) | 226100 (100%) |
| Characteristic Path Lengths | 3.556 | 3.556 |
| Average Number of Neighbours | 4.454 | 4.454 |
| Number of Nodes | 476 | 476 |
| Network Density | 0.009 | 0.009 |
| Isolated Nodes | 0 | 0 |
| Number of Self Loops | 0 | 0 |
| Multi-edge Node Pairs | 16 | 0 |
| Analysis Time (sec) | 0.071 | 0.018 |

The significant Transcription Factors in the inter-regulatory TFTG network were identified using Cytoscape plugin, CytoHubba. It was used to find the most essential modules and top 10 ranked nodes in the entire network. The 12 scoring methods used by CytoHubba to determine the critical network modules and top-ranked TFs/TGs in the inter-regulatory network include Betweenness centrality, BottleNeck, Closeness centrality, Clustering Coefficient, Degree centrality, EcCentricity, Edge Percolating Coefficient (EPC), Maximal Clique Centraity (MCC), Density of Maximum Neighbourhood Component (DMNC), Maximum Neighbourhood Component (MNC), Radiality and Stress centrality (Figure 7). The top 10 nodes in the network were determined by each scoring method respectively.

Based on their frequency of occurrence in each of the12 scoring methods, the top 5 Transcription Factors are represented in Figure 2A and 2B. The analysis of the TFTG network identified 9 transcription factors namely Mef2c, Isl1, Fosl2, Rxra, Jun, Pparg, Ascl1, Onecut2 and Gata1 to be significantly enriched; out of which Mef2c and Isl1 were the two most critical transcription factors having the highest frequency of occurrence in the *L. major* infection (Figure 8 E).

## Discussion

### Macrophage cell clustering dynamics changes with time point of infection

As compared to control samples 6hr sample showed similar number of clusters although the dynamics of clusters shifted in t-SNE plot with changes in number of cells in individual clusters. At 12hr, cluster size and number increased with cell distribution among 12 clusters. At 18hr cluster number changed to 10, we do not have any apparent reason for this change; we predict that the macrophage plasticity was modulated by parasite so that number of cells in a particular cluster increases so the number of phenotypic macrophages decreases.

### Enrichment of M1 and M2 macrophages unveiled IL10 and IL12 reciprocity in *L.major* infection

Two main phenotypes of macrophages are M1 and M2 which offers inflammatory and regulatory roles respectively in leishmaniasis. They differ mainly in production of chemokines and cytokines, transcription factors and specific markers which identifies their function (Tomiotto-Pellissier et al., 2018). IFNγ is involved in classical activation of macrophage to M1 phenotye (Orecchioni et al., 2019), CD80 is an expression marker of M1 macrophages (Atri et al., 2018) whereas IL10 promotes alternative activation of macrophage to M2 phenotype commonly in all M2 subtypes (Lopes et al., 2016) (Tomiotto-Pellissier et al., 2018). Hence, we used them to identify overall M1 and M2 subtypes at all time points and observed the dominance of M2 over M1 population this maybe because of amastigotes driven modulation of macrophage plasticity at with time of infection. Identification of M2 subsets was one of our novel findings as categorization of M2 subtypes was possible through sophisticated parameter provided by scATAC-seq. Typically, log2 values harnessed are set as more than 0.5 (Merrill et al., 2022), even so we set threshold as 1 to ensure capturing of transitioned M1 and M2 macrophages and to avoid identification of mixed population. Enrichment of M2a suggested the induction and over expression of arginase and IL10 and enrichment of M2d suggested increase in IL10 levels and decreased IL12 expression.

### IL10 expression is more prevalent in *L.major* infection

From M1 and M2 expression paradigm we wanted to identify the populations which are were potent in expressing IL10 and IL12. Hence we filtered the population which had expressed IL10 more than IL12 and vice versa to avoid overlap of population. Sudden increase of IL12 producing macrophages at 12hr post infection suggested that macrophages were trying to combat infection nonetheless their population subsided at 18hr due to overexpression of IL10 by majority of population.

### Reciprocal expression of IL10 and IL12 occurs post 6hrs of infection

Through our previous work we had reported the modulation of IL10 and IL12 by NFAT5 and SHP-1 in *L.major* infection models through systems based discrete mathematical models. NFAT5 being transcription factor and chromatin remodelling inducer, it remodels nucleosome1 of IL12b gene and nucleosome2 of IL10 promoter region to enable parasite killing. Later, as a mechanism to combat infection and regulate pro-inflammatory cytokine, *Leishmania* modulates host machinery to activate SHP-1 a phosphatase; to inactivate NFAT5 and therefore the chromatin architecture changes entirely which activates IL10 gene (Khandibharad and Singh, 2022a). Our systems based findings were validated when we observed expressional changes in IL10 and IL12 expression post *L.major* infection from 1hr post infection till 48hr with major changes occurring at 6hr, 12hr and 18hr (Khandibharad and Singh, 2022b).

The expression based findings from scATAC-seq data corroborates with our data. We checked co-expression of parasite eliminating group which included IL12, NFAT5 and iNOS genes comparatively with parasite survival favouring group which included IL10, SHP-1 and Arg1. iNOS and Arginase expression were also considered for co-expression as ultimately production of nitric oxide defines the parasite survival fate. At 6hr we observed parasite elimination macrophages dominating with decline in the population at 12hr and switch in macrophage population from parasite eliminating to parasite survival favouring at 18hr. Thus this study unveiled the importance of time dependant cellular changes which leads to leishmaniasis.

### Sleepy macrophages acts as a cellular switch of macrophage phenotypes by channelling cell plasticity

Systematically we started analysing all the samples for their expression pattern, we checked the expression of housekeeping genes ActB and Gapdh, unexpectedly we observed cluster 3 and cluster 6 from 6hr sample does not express housekeeping genes. To our astonishment, these cells did not express IL12b, IL10, Ptpn6 (SHP-1) and Nfat5 although the motif z-score for NFAT5 was highest in these two clusters than across any other cluster in the same sample. These unique clusters were absent in other samples. Since these clusters consisted overall 28% of the population we got intrigued about the set of genes these clusters express, transcription factors associated with these genes, pathways which are active in these clusters and most importantly crucial components in these clusters which are making them unique to such behaviour. As these macrophages were not expressing housekeeping genes, genes which we were studying and any marker genes which macrophages normally express, we named them as “sleepy macrophages” to describe their dormancy. We predict that amastigotes inside these macrophages are activating gene sets which may be conferring to their uniqueness.

### Principle component analysis revealed Cluster 3 and its genes are critical in sleepy macrophages

The first observation which we made was the genes which are upregulated in Cluster 3 and 6 were downregulated in Cluster 1, 2, 4 and 5. Hence to identify which cluster is essential we did the PCA and observed that 92. 1% information is retained by first four clusters. From variable plot we can also say that Cluster 3 is negatively correlated with other clusters with high cos2 value. As an individual PC Custer3 is highly correlated with PC1 and PC2 and Cluster 6 had lesser correlation suggesting its low contribution in the data set. We distributed the PCs based on significance of their expression and categorized 814 genes in group of 5 for all clusters of which genes having p-value of more than 0.0001 were identified and they showed low redundancy distribution. Hence from this analysis we focused our studies on Cluster 3 to identify behaviour of sleepy macrophages.

### Sleepy macrophages acts as transient state to prepare macrophages to combat *L.major* infection

Using Gene Set Enrichment Analysis (GSEA) we were able to analyse the defined set of genes from 6hr sample by measuring statistically significant and concordant changes between phenotypes (diseased and normal). GSEA was used to evaluate the relevance of numerous aberrations in gene expression and cellular transcriptional responses using single cell ATAC gene expression dataset for *Mus musculus* macrophage. Using the gene expression profiles of infected macrophages, GCT, GMT, and CLS files were prepared as input files for GSEA analysis (Supplementary file 2 (S9 and S9)). Metric used for ranking the genes in the dataset was Diff_of_Classes and chip platform provided for the analysis was Mouse_NCBI_Gene_ID_MSigDB.v2022.1.Mm.chip. The results obtained showed that the genes belonging to cluster 3 and 6 were positively correlated with the diseased phenotype or state.

We analyzed top 10 genes from heatmap by excluding pseudogenes and observed Pcdh11x was enriched in cluster 3 which is responsible for cell recognition and activation of PI3k/Akt signaling (Wei et al., 2022), Taf2 is critical in formation of RNA polymerase II initiation complex (Alpern et al., 2014), TMEM117 primarily functions in Endoplasmic reticulum (ER) stress mediated mitochondrial apoptotic pathway (Maruyama et al., 2023), Gabrg1 has a role in chloride channel activity although we could not find its relevance in leishmaniasis or macrophages, Tox is a DNA binding protein associated with control of chromatin structure majorly in activation of T-cells (Yu and Li, 2015), Fastkd3 is an unusual RNA-binding protein that critically regulates mitochondrial RNA metabolism (Boehm et al., 2016), *L.major* infected mice model has been reported to show significant reduction in parasite burden in lymph nodes, spleen and liver when Map2 was activiity was enhanced in vivo (Le Pape, 2008), Erap2 acts as protease to cleave antigens and form peptides which are presented by MHCI in canine leishmaniasis (Pedersen et al., 2016), Zfp606 has not been reported to have any role, Vmn2r70 is predicted to have a role in enabling G protein-coupled receptor activity. From the expression set we could infer that sleepy macrophages have a stressful environment and is highly undergoing epigenetic ad transcriptional changes to *L.major* infection and is preparing it to combat infection by itself and also to act as an antigen presenting cell to other immune cells.

### NFAT5 controls cellular signaling and epigenetically regulates sleepy macrophages as a response to *L.major* infection

Mef2c has been demonstrated to be associated with enrichment of classical activated macrophages that is M1 and promotes pro-inflammatory cytokine production in leishmaniasis (Dirkx et al., 2022), it was observed from patient samples that synthesis of Mef2c is associated with NFAT5 canonical pathway which also in turn regulates NFAT5 mediated immune response, TNFα is the upstream regulator of MEF2C (Salih et al., 2017). Another transcription factor which was enriched through TFTG network was Isl1, although much is not known about the role of this gene in leishmaniasis nonetheless, it consists of LIM domain is essential for regenerating T cell in spleen in leishmaniasis (Golub et al., 2018). From pathway enrichment analysis of all 814 genes and 51 transcription factors we observed NFAT5 signaling, gene transcription, MAPK signaling, TGFβ signaling were enhanced. From this analysis we could infer that NFAT5 mediated signaling and immune response is prevailing in sleepy macrophages.

## Conclusion

Our findings identify sleepy macrophages which possess a state adaptation to *L.major* infection. These cells boast pro-inflammatory cytokine secretion by promoting chromatin remodeling and RNA regulation through transcription factors. One such transcription fator which we have highlighted from our findings is NFAT5 that dictates IL10 and IL12 reciprocal regulation in *L.major* infection. Often NFAT5 gets downregulated by *L.major* induced SHP-1 mediated dephosphorylation of its auxillary export domain. It may be upright to target NFAT5 so as to direct adequate parasite elimination response. We presume that inhibition of SHP-1 may prevent NFAT5 inhibition and supress parasite survival. Our previous work has already underlined the proposition in which peptides will be used to inhibit SHP-1. In future, we will inspect the efficacy of peptides on macrophage population in *L.major* infection.

## References

1. Afrin F, Khan I, Hemeg HA. 2019. Leishmania-Host Interactions-An Epigenetic Paradigm. Front Immunol 10:492. doi:10.3389/fimmu.2019.00492

2. Almeida L, Silva JA, Andrade VM, Machado P, Jamieson SE, Carvalho EM, Blackwell JM, Castellucci LC. 2017. Analysis of expression of FLI1 and MMP1 in American cutaneous leishmaniasis caused by Leishmania braziliensis infection. Infect Genet Evol J Mol Epidemiol Evol Genet Infect Dis 49:212–220. doi:10.1016/j.meegid.2017.01.018

3. Alpern D, Langer D, Ballester B, Le Gras S, Romier C, Mengus G, Davidson I. 2014. TAF4, a subunit of transcription factor II D, directs promoter occupancy of nuclear receptor HNF4A during post-natal hepatocyte differentiation. Elife 3:e03613. doi:10.7554/eLife.03613

4. Andersen L, Corazon SSS, Stigsdotter UKK. 2021. Nature Exposure and Its Effects on Immune System Functioning: A Systematic Review. Int J Environ Res Public Health 18. doi:10.3390/ijerph18041416

5. Atri C, Guerfali FZ, Laouini D. 2018. Role of Human Macrophage Polarization in Inflammation during Infectious Diseases. Int J Mol Sci 19. doi:10.3390/ijms19061801

6. Baek S, Lee I. 2020. Single-cell ATAC sequencing analysis: From data preprocessing to hypothesis generation. Comput Struct Biotechnol J 18:1429–1439. doi:10.1016/j.csbj.2020.06.012

7. Baharia RK, Tandon R, Sahasrabuddhe AA, Sundar S, Dube A. 2021. Correction: Nucleosomal Histone Proteins of L. donovani: A Combination of Recombinant H2A, H2B, H3 and H4 Proteins Were Highly Immunogenic and Offered Optimum Prophylactic Efficacy against Leishmania Challenge in Hamsters. PLoS One. doi:10.1371/journal.pone.0252177

8. Baker RE, Mahmud AS, Miller IF, Rajeev M, Rasambainarivo F, Rice BL, Takahashi S, Tatem AJ, Wagner CE, Wang L-F, Wesolowski A, Metcalf CJE. 2022. Infectious disease in an era of global change. Nat Rev Microbiol 20:193–205. doi:10.1038/s41579-021-00639-z

9. Baker SM, Rogerson C, Hayes A, Sharrocks AD, Rattray M. 2019. Classifying cells with Scasat, a single-cell ATAC-seq analysis tool. Nucleic Acids Res 47:e10. doi:10.1093/nar/gky950

10. Boehm E, Zornoza M, Jourdain AA, Delmiro Magdalena A, García-Consuegra I, Torres Merino R, Orduña A, Martín MA, Martinou J-C, De la Fuente MA, Simarro M. 2016. Role of FAST Kinase Domains 3 (FASTKD3) in Post-transcriptional Regulation of Mitochondrial Gene Expression. J Biol Chem 291:25877–25887. doi:10.1074/jbc.M116.730291

11. Chen H, Lareau C, Andreani T, Vinyard ME, Garcia SP, Clement K, Andrade-Navarro MA, Buenrostro JD, Pinello L. 2019. Assessment of computational methods for the analysis of single-cell ATAC-seq data. Genome Biol 20:241. doi:10.1186/s13059-019-1854-5

12. Curtin JM, Aronson NE. 2021. Leishmaniasis in the United States: Emerging Issues in a Region of Low Endemicity. Microorganisms 9. doi:10.3390/microorganisms9030578

13. Dirkx L, Hendrickx S, Merlot M, Bulté D, Starick M, Elst J, Bafica A, Ebo DG, Maes L, Van Weyenbergh J, Caljon G. 2022. Long-term hematopoietic stem cells as a parasite niche during treatment failure in visceral leishmaniasis. Commun Biol 5:626. doi:10.1038/s42003-022-03591-7

14. Fang R, Preissl S, Li Y, Hou X, Lucero J, Wang X, Motamedi A, Shiau AK, Zhou X, Xie F, Mukamel EA, Zhang K, Zhang Y, Behrens MM, Ecker JR, Ren B. 2021. Comprehensive analysis of single cell ATAC-seq data with SnapATAC. Nat Commun 12:1337. doi:10.1038/s41467-021-21583-9

15. Fukushima A. 2013. DiffCorr: an R package to analyze and visualize differential correlations in biological networks. Gene 518:209–214. doi:10.1016/j.gene.2012.11.028

16. Golub R, Tan J, Watanabe T, Brendolan A. 2018. Origin and Immunological Functions of Spleen Stromal Cells. Trends Immunol 39:503–514. doi:10.1016/j.it.2018.02.007

17. J B, M BM, Chanda K. 2021. An Overview on the Therapeutics of Neglected Infectious Diseases-Leishmaniasis and Chagas Diseases. Front Chem 9:622286. doi:10.3389/fchem.2021.622286

18. Jia G, Preussner J, Chen X, Guenther S, Yuan X, Yekelchyk M, Kuenne C, Looso M, Zhou Y, Teichmann S, Braun T. 2018. Single cell RNA-seq and ATAC-seq analysis of cardiac progenitor cell transition states and lineage settlement. Nat Commun 9:4877. doi:10.1038/s41467-018-07307-6

19. Kamhawi S, Serafim TD. 2020. Leishmania: A Maestro in Epigenetic Manipulation of Macrophage Inflammasomes. Trends Parasitol 36:498–501. doi:10.1016/j.pt.2020.04.008

20. Khandibharad S, Nimsarkar P, Singh S. 2022. Mechanobiology of immune cells: Messengers, receivers and followers in leishmaniasis aiding synthetic devices. Curr Res Immunol 3:186–198. doi:10.1016/j.crimmu.2022.08.007

21. Khandibharad S, Singh S. 2022a. Computational System Level Approaches for Discerning Reciprocal Regulation of IL10 and IL12 in Leishmaniasis. Front Genet 12:1–14. doi:10.3389/fgene.2021.784664

22. Khandibharad S, Singh S. 2022b. Artificial intelligence channelizing protein-peptide interactions pipeline for host-parasite paradigm in IL-10 and IL-12 reciprocity by SHP-1. Biochim Biophys acta Mol basis Dis 1868:166466. doi:10.1016/j.bbadis.2022.166466

23. Le Pape P. 2008. Development of new antileishmanial drugs--current knowledge and future prospects. J Enzyme Inhib Med Chem 23:708–718. doi:10.1080/14756360802208137

24. Lê S, Josse J, Husson F. 2008. FactoMineR: An R Package for Multivariate Analysis. J Stat Softw 25:1–18. doi:10.18637/jss.v025.i01

25. Liu D, Uzonna JE. 2012. The early interaction of Leishmania with macrophages and dendritic cells and its influence on the host immune response. Front Cell Infect Microbiol 2:83. doi:10.3389/fcimb.2012.00083

26. Lopes RL, Borges TJ, Zanin RF, Bonorino C. 2016. IL-10 is required for polarization of macrophages to M2-like phenotype by mycobacterial DnaK (heat shock protein 70). Cytokine 85:123–129. doi:10.1016/j.cyto.2016.06.018

27. Loría-Cervera EN, Andrade-Narvaez F. 2020. The role of monocytes/macrophages in Leishmania infection: A glance at the human response. Acta Trop 207:105456. doi:10.1016/j.actatropica.2020.105456

28. Maruyama R, Kiyohara Y, Kudo Y, Sugiyama T. 2023. Effects of the anti-inflammatory drug celecoxib on cell death signaling in human colon cancer. Naunyn Schmiedebergs Arch Pharmacol. doi:10.1007/s00210-023-02399-4

29. Masina S, M Gicheru M, Demotz SO, Fasel NJ. 2003. Protection against cutaneous leishmaniasis in outbred vervet monkeys using a recombinant histone H1 antigen. J Infect Dis 188:1250–1257. doi:10.1086/378677

30. Merrill CB, Montgomery AB, Pabon MA, Shabalin AA, Rodan AR, Rothenfluh A. 2022. Harnessing changes in open chromatin determined by ATAC-seq to generate insulin-responsive reporter constructs. BMC Genomics 23:399. doi:10.1186/s12864-022-08637-y

31. Mukherjee S, Mukherjee B, Mukhopadhyay R, Naskar K, Sundar S, Dujardin J-C, Roy S. 2014. Imipramine exploits histone deacetylase 11 to increase the IL-12/IL-10 ratio in macrophages infected with antimony-resistant Leishmania donovani and clears organ parasites in experimental infection. J Immunol 193:4083–4094. doi:10.4049/jimmunol.1400710

32. Mulqueen RM, Pokholok D, O’Connell BL, Thornton CA, Zhang F, O’Roak BJ, Link J, Yardımcı GG, Sears RC, Steemers FJ, Adey AC. 2021. High-content single-cell combinatorial indexing. Nat Biotechnol 39:1574–1580. doi:10.1038/s41587-021-00962-z

33. Orecchioni M, Ghosheh Y, Pramod AB, Ley K. 2019. Macrophage Polarization: Different Gene Signatures in M1(LPS+) vs. Classically and M2(LPS-) vs. Alternatively Activated Macrophages. Front Immunol 10:1084. doi:10.3389/fimmu.2019.01084

34. Pedersen NC, Dhanota JK, Liu H. 2016. Polymorphisms in ERAP1 and ERAP2 are shared by Caninae and segregate within and between random-and pure-breeds of dogs. Vet Immunol Immunopathol 179:46–57. doi:10.1016/j.vetimm.2016.08.006

35. Pott S, Lieb JD. 2015. Single-cell ATAC-seq: strength in numbers. Genome Biol 16:172. doi:10.1186/s13059-015-0737-7

36. Rai V, Quang DX, Erdos MR, Cusanovich DA, Daza RM, Narisu N, Zou LS, Didion JP, Guan Y, Shendure J, Parker SCJ, Collins FS. 2020. Single-cell ATAC-Seq in human pancreatic islets and deep learning upscaling of rare cells reveals cell-specific type 2 diabetes regulatory signatures. Mol Metab 32:109–121. doi:10.1016/j.molmet.2019.12.006

37. Salih MAM, Fakiola M, Lyons PA, Younis BM, Musa AM, Elhassan AM, Anderson D, Syn G, Ibrahim ME, Blackwell JM, Mohamed HS. 2017. Expression profiling of Sudanese visceral leishmaniasis patients pre-and post-treatment with sodium stibogluconate. Parasite Immunol 39. doi:10.1111/pim.12431

38. Subramanian A, Tamayo P, Mootha VK, Mukherjee S, Ebert BL, Gillette MA, Paulovich A, Pomeroy SL, Golub TR, Lander ES, Mesirov JP. 2005. Gene set enrichment analysis: a knowledge-based approach for interpreting genome-wide expression profiles. Proc Natl Acad Sci U S A 102:15545–15550. doi:10.1073/pnas.0506580102

39. Sundar S, Singh B. 2018. Emerging therapeutic targets for treatment of leishmaniasis. Expert Opin Ther Targets 22:467–486. doi:10.1080/14728222.2018.1472241

40. Tomiotto-Pellissier F, Bortoleti BT da S, Assolini JP, Gonçalves MD, Carloto ACM, Miranda-Sapla MM, Conchon-Costa I, Bordignon J, Pavanelli WR. 2018. Macrophage Polarization in Leishmaniasis: Broadening Horizons. Front Immunol 9:2529. doi:10.3389/fimmu.2018.02529

41. Wei X, Yuan Y, Yang Q. 2022. SNHG22 promotes migration and invasion of trophoblasts via miR-128-3p/PCDH11X axis and activates PI3K/Akt signaling pathway. Clinics (Sao Paulo) 77:100055. doi:10.1016/j.clinsp.2022.100055

42. Wu X, Lu M, Yun D, Gao S, Chen S, Hu L, Wu Y, Wang X, Duan E, Cheng CY, Sun F. 2022. Single-cell ATAC-Seq reveals cell type-specific transcriptional regulation and unique chromatin accessibility in human spermatogenesis. Hum Mol Genet 31:321–333. doi:10.1093/hmg/ddab006

43. Xu W, Wen Y, Liang Y, Xu Q, Wang X, Jin W, Chen X. 2021. A plate-based single-cell ATAC-seq workflow for fast and robust profiling of chromatin accessibility. Nat Protoc 16:4084–4107. doi:10.1038/s41596-021-00583-5

44. Yu X, Li Z. 2015. TOX gene: a novel target for human cancer gene therapy. Am J Cancer Res 5:3516–3524.

45. Zhu T, Liao K, Zhou R, Xia C, Xie W. 2020. ATAC-seq with unique molecular identifiers improves quantification and footprinting. Commun Biol 3:675. doi:10.1038/s42003-020-01403-4

46. Zito A, Lualdi M, Granata P, Cocciadiferro D, Novelli A, Alberio T, Casalone R, Fasano M. 2021. Gene Set Enrichment Analysis of Interaction Networks Weighted by Node Centrality. Front Genet 12:577623. doi:10.3389/fgene.2021.577623

